# Preclinical evaluation of pharmacological inhibition of SIRT1 on the growth of tumoral and metastatic granulosa cells

**DOI:** 10.1101/2024.07.02.601506

**Authors:** Victoria Cluzet, Eloïse Airaud, Marie M Devillers, Florence Petit, Alexandra Leary, Alice Pierre, Haojian Li, Chi-Ping Day, Urbain Weyemi, Stéphanie Chauvin, Céline J Guigon

## Abstract

**Background:** Clinical management of patients with recurrent ovarian granulosa cell tumor (GCT) remains poor. Sirtuin-1 (SIRT1), a deacetylase enzyme involved in the regulation of tumor growth and metastasis, may represent a therapeutic target due to the availability of selective pharmacological inhibitors with minimal toxicity.

**Methods:** We assessed the possible overexpression of SIRT1 during tumorigenesis by Western blot and immunohistochemistry. We tested the effects of SIRT1 inhibition by EX-527 on growth, proliferation, death, migration and gene expression by RNA sequencing and RT-qPCR *in vitro* on three GCT cell lines (AT29, KGN, COV434). Tumor growth in response to EX-527 treatment was examined in nude mice carrying subcutaneous GCT cell grafts using an electronic caliper and in GCT of AT83 mice by 3D ultrasound imaging system.

**Results:** SIRT1 abundance increased during tumorigenesis. *In vitro* treatment with EX-527 efficiently reduced cell growth, either by inducing apoptosis or by inhibiting proliferation. EX-527 induced alterations in pathways driven by mTOR, Myc and E2F, and in pathways controlling cell metabolism and oxidative stress. The administration of this treatment for 4 weeks efficiently reduced tumor progression *in vivo*.

**Conclusions:** Our study reveals a new therapeutic potential of SIRT1 targeting as a treatment option for patients with recurrent GCT.

## Introduction

Granulosa cell tumors (GCT) of the ovary represent about 7% of ovarian cancers. They present two distinct clinical presentations, the juvenile form which occurs mainly in childhood and adolescence, and the adult form which occurs mainly in peri/post-menopausal women [1]. Although this type of tumor is often diagnosed at an early stage of progression, it has a high risk of late recurrence with metastases spreading to the peritoneal cavity and liver [2]. Additionally, patients with GCT frequently present with endocrine alterations, including elevated estradiol levels that can lead to menorrhagia, irregular menstruation, precocious puberty, and possibly promote the progression of GCT [1,3,4]. Chemotherapy and hormonotherapy with aromatase inhibitors globally give unsatisfactory outcomes, as about 80% of patients with advanced tumors eventually die [5].

The etiology of this form of cancer is still elusive but converging evidence suggests that Forkhead box L2 transcription factor (FOXL2) plays an essential role in tumorigenesis. Indeed, 97% of women with adult GCT have a somatic mutation (referred to as c.C402G (p.Cys134Trp) mutation) [6]. Mice carrying the human C134W *FOXL2* mutation develop GCT with full penetrance by 18 months of age [7]. Although this mutation is not present in juvenile GCT, FOXL2 expression is absent or low in ∼50% of patients with aggressive tumors [8]. Additional somatic mutations have been observed in primary and recurrent GCT, including in the *TERT* (telomerase) promoter, *KMT2D* (Lysine methyltransferase 2D) and *TP53* [9–12]. We have previously reported the variations in the expression and types of estrogen receptors (ERs) present in the GCT that may underlie the variable effectiveness of aromatase inhibitors in patients [3,13]. These findings illustrate tumor heterogeneity and the difficulty of finding effective treatments for most patients, highlighting the need for personalized medicine.

Sirtuin-1 (SIRT1) belongs to the family of NAD+-dependent enzymes that regulate fundamental biological functions ranging from genomic stability and lifespan to energy metabolism and tumorigenesis. This enzyme deacetylates a number of histone and non-histone substrates such as P53 and E2F to regulate their action [14]. It negatively regulates the expression or activity of FOXL2 and p53 in GCT cell lines [15,16], and acts in concert with ERα in breast cancer to mediate estradiol oncogenic actions [17]. SIRT1 is often expressed in GCT [18], where it may participate to cell proliferation and survival [16,18–20]. It is overexpressed in many cancers and its pharmacological inhibition can slow tumor growth and progression in animal models [14,21,22]. These results make SIRT1 a potential therapeutic target for GCT. Among the selective SIRT1 inhibitors, EX-527 (6-chloro-2,3,4,9-tetrahydro-1H-carbazole-1-carboxamide), is one of the few compounds for which initial mechanistic data are available, combining high potency with significant isoform selectivity [23,24]. EX-527 inhibits SIRT1 ∼100-fold more potently than SIRT2 and SIRT3 and has no effect on SIRT5 deacetylation activity [25]. Additionally, in clinical trials, this SIRT1 inhibitor has been shown to be well tolerated and safe in healthy subjects [26]. Taken together, these observations suggest that suppression of SIRT1 enzymatic activity by EX-527 could be an attractive therapeutic option in patients with GCT, but sufficient evidence is lacking.

The aim of this study was to investigate the potential benefit of targeting SIRT1 with EX-527 on tumor growth and progression. We first sought for possible overexpression of SIRT1 in GCT of patients and mice. We then evaluated the efficacy of EX-527 treatment *in vitro* using GCT cell lines. We finally extended our investigations to preclinical mouse models to validate our *in vitro* results on the ability of EX-527 to slow GCT growth.

## Materials and Methods

### Granulosa cells and GCT from patients

Luteinized granulosa cells (GC) were obtained from patients undergoing *in vitro* fertilization at Antoine Béclère Hospital (France) and were purified as previously described [3]. All women met the inclusion criteria listed in [3]. Informed consent was obtained from patients, and these studies had the approval of the internal institutional review board of Antoine Béclère Hospital under an Institutional Review Board-exempt protocol. The investigation received the approval of internal institutional review board, IRB Blefco-IORG0010582. Seven adult primary GCT retrieved at an advanced stage were used [3], with the approval of the internal institutional review board from the Institut Gustave Roussy under an Institutional Review Board-exempt protocol. The study was performed in accordance with the Declaration of Helsinki for Medical Research involving Human Subjects (2013 revision).

### Transgenic mice with GCT

AT83 mice were obtained by targeted oncogenesis of the anti-Müllerian hormone promoter-SV40 oncogene construct [27,28]. They were maintained under controlled conditions (12 hours light/12 hours dark) with food (Scientific Animal Food and Engineering, A03-10) and water available *ad libitum*. For ovary collection, mice were anesthetized with a mix of ketamine (Imalgene® 1000) and xylazine (Rompun® 2%). After cervical dislocation, ovaries were collected, weighed, and either frozen in liquid nitrogen and stored at −80°C for protein extraction, or fixed in 4% paraformaldehyde (PFA) for immunohistochemistry. For *in vivo* follow-up of tumor growth using ultrasound imaging (see below), AT83 mice > 12 months with palpable tumors were included in the studies. Experiments were performed in accordance with standard ethics guidelines and were approved by Institutional Animal care and Use committee of the Université Paris Cité and by the French Ministry of Agriculture (agreement #02193.02).

### GCT cell lines

The AT29 cell line was established from a primary GCT of the AT transgenic mouse and it retains a number of specific granulosa cell markers such as SF-1, WT-1, AMHR2 and FOXL2 [3,27,28]. It was generously provided by Dr di Clemente (Sorbonne Université, INSERM, Centre de recherche Saint-Antoine). Only cells at an early passage (<30) were used. The KGN cells line was established from an adult patient with recurrent, metastasized GCT in the pelvic region [29]. It harbors the *FOXL2 C402G* mutation described in most adult GCT, in addition to the *TERT* promoter and *KMT2D* mutations [10,11]. It was purchased from the RIKEN BioResource Center (RBRC-RCB1154, RIKEN Cell Bank, Ibaraki, Japan) after approval by Dr Yoshiro Nishi and Dr Toshihiko Yanase. COV434 cells, derived from a primary tumor of a 27-year-old patient [30], display the wild-type *FOXL2* gene but they do not express it, as observed in 50% of aggressive GCT of the juvenile isotype [31]. These cells were purchased from ECACC (#07071909, Sigma-Aldrich). Recent studies questioned the fact that COV434 cells are of GCT origin and proposed instead that they represent small cell carcinoma of the ovary, hypercalcemic type (SCCOHT), based notably on the absence of *SMARCA2* and *SMARCA4* mRNA and protein [32,33]. In contrast with these findings, we found *SMARCA2* and *SMARCA4* mRNA in COV434 cells, in addition to that of AMH which is a well-known marker of granulosa cells and considered as a marker of GCT (Supplementary Table 1). Taken together, this data supports the idea that the COV434 cell line used in the present study is of GCT origin. GCT cell lines were routinely maintained at 37°C with 5% CO_2_ in complete medium consisting of Dulbecco’s modified Eagle medium: nutrient mixture F-12 (DMEM/F12) (Gibco) containing 10% fetal bovine serum (FBS) and 0.5% penicillin/streptomycin.

### SIRT1 immunohistochemistry in ovaries and GCT of mice and women

Immunohistochemistry on mouse and human paraffin-embedded tissues were performed as previously described, with some modifications [34]. Briefly, rehydrated ovarian or GCT sections were heated to 98°C in 0.05% citraconic anhydride, pH 7.4 (Sigma-Aldrich Corp.) for 45 minutes and then blocked in 10% FBS/1% bovine serum albumin (BSA) in phosphate buffered saline (PBS) for 1 hour. After washing in PBS, slides of human ovaries and GCT were incubated overnight at 4°C, while those of mice were incubated for 30 minutes with the mouse monoclonal SIRT1 antibody (dilution 1/200) (Table 1). After washing in PBS, slides were incubated with anti-mouse secondary antibodies (Dako MOM kit, PK-2200, Vector Laboratories, Burlingame, CA; SK-4100) for 1 hour. Slides were then incubated in 3,3′-diaminobenzidine (DAB substrate kit for peroxidase; Vector Laboratories), and after staining development, they were counterstained in Hematoxylin, rinsed and mounted in Eukitt (Sigma).

**Table 1:** List of antibodies used for immunohistochemistry and Western blot.

| Target | Provider | Reference | Isotype | Dilution |
| --- | --- | --- | --- | --- |
| Uncleaved Caspase-8 | Cell signaling | #4927 | rabbit polyclonal | 1/1000 |
| Cleaved Caspase-8 | Cell signaling | #8592 | rabbit polyclonal | 1/1000 |
| Cyclin D2 | Santa Cruz | sc-181 | rabbit polyclonal | 1/200 |
| GAPDH | Santa Cruz | sc-25778 | rabbit polyclonal | 1/4000 |
| SIRT1 | Cell signaling | #8469 | mouse monoclonal | 1/4000 |
| Vinculin | Sigma-Aldrich | V9131 | mouse monoclonal | 1/120000 |

### Western blots

Single frozen human and mouse GCT, human luteinized GC, WT mouse ovaries and AT29 cells allografts were lysed with TissueLyser (Qiagen) in Frackelton buffer, as previously described [27]. Protein samples (30 μg) were loaded and separated by sodium dodecyl sulfate–polyacrylamide gel electrophoresis, and then electrotransferred to nitrocellulose membranes. Membranes were then incubated in 10% milk for 60 minutes and incubated overnight with primary antibodies (Table 1) diluted in 5% milk or BSA at 4°C. The following day, the membranes were incubated with anti-rabbit or anti-mouse HRP-linked IgG antibodies (dilution: 1/3000) depending on primary antibodies, for 90 minutes. Proteins were detected by chemiluminescence with Pierce^TM^ ECL Plus Western Blotting substrate (#32132 Thermo Scientific) using a CCD camera (LAS 4000, FujiFilm or AI600, GE Healthcare).

### SIRT1 activity in GCT cell lines

AT29, KGN and COV434 cells were seeded at a density of 3×10^6^ cells in 100×15mm Petri dishes in complete medium. At confluence, cells were lysed with the lysis buffer provided by the SIRT1 Activity Assay Kit (#ab156065, Abcam^®^) (10mM Tris, HCl (pH 7.5), 10mM NaCl, 15mM MgCl2, 250mM Sucrose, 0.5% NP-40, 0.1mM EGTA) and sonicated four times during 5 seconds. After a 10 minute-centrifugation at 15 000 rpm at 4°C, SIRT1 antibody (#8469S, Cell Signaling Technology^®^) was added at a dilution of 1/62 to supernatants and incubated overnight under constant agitation at 4°C. The following day, supernatants were incubated with 30 µL of protein G Sepharose™ 4 Fast Flow (#17-0618-01, GE Healthcare) under agitation for 4 hours at 4°C. The precipitate was then washed three times with a 30 second-centrifugation at 12 000 *g* at 4°C, and the pellet resuspended in 25 µL of distilled water. SIRT1 activity was then measured according to the manufacturer’s instructions. Briefly, the precipitates were incubated with 25 µL of a reagent mix including assay buffer, coenzyme nicotinamide adenine dinucleotide (NAD+), developer, fluoro-substrate peptide and of one of the following treatments: 1/100 dilution of dimethyl-sulfoxide (DMSO), 20 µM or 50 µM of EX-527 in black 96-well plates. Positive controls were prepared with recombinant SIRT1 incubated with EX-527 at 20 and 50 µM. Negative controls were performed using recombinant SIRT1 and the reagent mix without NAD+. SIRT1 activity was measured by reading fluorescence at an emitted light of 450 nm every 2 minutes during 2 hours, using FlexStation 3 (Molecular Devices).

### GCT cell line growth by MTT assay

For MTT (3-(4,5-dimethylthiazole-2-yl)-2,5-diphenyl tetrazolium bromide) assay, AT29, KGN and COV434 cells were seeded at a density of 5×10^3^ cells per well for AT29 and 4×10^4^ for KGN and COV434 in 24-well plates in complete medium. Twenty-four hours later, they were treated with EX-527 (#2780, Tocris, 50 μM) in complete medium. After 0, 24, 48 and 72 hours of treatment, cells were incubated with MTT (1 mg/mL in Dulbecco’s phosphate-buffered saline [DPBS]) for 2 hours, and then lysed in DMSO. The absorbance was read at 575 nm with FlexStation3 (Molecular Devices). The experiments were run at least three times.

### GCT cell line viability and apoptosis with ApoTox-Glo^TM^ Triplex assay

Cell viability and apoptosis was evaluated by the ApoTox-Glo^TM^ Triplex assay system (#G6320, Promega) in line with the manufacturer’s instructions. Cells were seeded at a density of 1×10^4^ in 96-well plates, with white wall and clear bottom in complete medium. Twenty-four hours later, cells received treatments with DMSO (CTR) or EX-527 (50 μM) for 24 hours in complete medium. Then, 20 μL of viability and cytotoxicity reagent (GF-AFC substrate for viable cells, and bis-AAF-R110 substrate for dead cells) were added in each well. Fluorescence intensities of substrates were measured using a microplate reader (FlexStation 3) at the following two wavelength sets: 400_Ex_/505_Em_ (Viability); 485_Ex_/520_Em_ (Cytotoxicity) at room temperature. 100 μL of Caspase-Glo reagents were then added into each well and the kinetic of luminescence intensity was measured using Flexstation3 for 3 hours at room temperature. For each sample, the maximal caspase-3/7 activity obtained during kinetic analysis was normalized to the ratio of the number of live cells to the number of dead cells.

### GCT cell line proliferation by ELISA assays

Measurement of BrdU incorporation was analyzed using the Cell Proliferation ELISA BrdU colorimetric test (#11647229001, Roche), in line with the manufacturer’s instruction. AT29 and KGN cells were seeded in complete medium at a density of 5×10^3^ cells per well, and COV434 cells at a density of 1×10^4^ cells, in 96-well plates with white wall and clear bottom and treated 24 hours later as for the ApoTox-GloTM assay. At the end of treatments, cells were incubated with the BrdU labeling solution diluted in DMEM/F12 medium without FBS, penicillin/streptomycin and phenol red (final concentration: 10 μM). Then, cells were fixed and marked with an anti-BrdU-peroxidase antibody for 90 minutes. BrdU incorporation was determined after addition of the peroxidase substrate by OD quantification at 370 nm and subtraction of the background measured at 492 nm by spectrometer (Flexstation 3, Molecular Devices).

### Oxidative stress

Oxidative stress was evaluated by the CellROX® Green Reagent (#C10444, Life Technologies) in line with the manufacturer’s instructions. AT29, KGN and COV434 cells were seeded in complete medium at a density of 1×10^5^, 1.5×10^5^ and 3×10^5^ cells, respectively, in 8-well Falcon® CultureSlides. Twenty-four hours later, cells received treatments with DMSO (CTR) or EX-527 (50 μM) for 24 hours in DMEM/F12 medium with 2% FBS and without phenol red. Then, cells were incubated with 5 µM of CellROX® Green Reagent for 30 minutes and were fixed with 4% PFA. Finally, slides were mounted with Fluoroshield™ containing DAPI (Sigma, F6057). Green (ROS) and blue (cell nuclei) fluorescence was observed with Nikon Eclipse 90i. The intensity of fluorescence in each treatment group was analyzed by ImageJ software (NIH, 1.64 version).

### RNA preparation from GCT cell lines

Cells were seeded in 6-well plates at a density of either 1,5×10^5^ (AT29 cells) or 2×10^5^ (COV434 and KGN cells) cells/mL in 10% FBS phenol red-free medium and treated with vehicle (CTR) or EX-527 at 50 μM for 24 hours. The experiment was replicated three times. A total of 18 samples for each cell line were lysed in RLT buffer from RNeasy mini kit (#74106, Qiagen) and RNAs were extracted following the manufacturer’s protocol. The quality and quantity of RNAs were estimated by Nanodrop (Thermo Scientific Nanodrop 2000).

### High-throughput RNA sequencing

For each cell line, three control and three EX-527-treated samples were obtained by pooling RNAs prepared from three different independent experiments in which three samples per condition were prepared. Total RNAs were qualified with AGILENT tapeStation 2200 (iGenSeq, ICM, France: https://igenseq.icm-institute.org). The preparation of mRNA library was realized following manufacturer’s recommendations (mRNA prep from ILLUMINA). Final samples pooled library prep were sequenced on ILLUMINA Novaseq 6000, corresponding to 2×44Millions of 100 bases reads per sample after demultiplexing. Quality of raw data was evaluated with FastQC. Poor quality sequences and adapters were trimmed or removed with fastp tool, with default parameters, to retain only good quality paired reads. Illumina DRAGEN bio-IT Platform (v3.8.4) was used for mapping on mm10 (i.e., 6 samples for AT29 cells) or hg38 (i.e., 6 samples for either COV434 or KGN cells) reference genome and quantification with gencode vM25 or gencode v37 annotation gtf file, respectively. Library orientation, library composition and coverage along transcripts were checked with Picard tools. Analysis was conducted with R software. Data were normalized with edgeR (v3.28.0) bioconductor packages prior to differential analysis with glm framework likelihood ratio test from edgeR package. Multiple hypothesis adjusted *P*-values were calculated with the Benjamini-Hochberg procedure to control FDR. Finally, enrichment analysis was conducted with clusterProfiler R package (v3.14.3) with Gene Set Enrichment Analysis and Hallmark and GeneOntology (GO). RNAseq data are available in a public, open access repository, under accession number GSE252773.

### Real-time PCR analysis

RNA from AT29, KGN and COV434 cell lines (700 ng) were reverse transcribed with the Superscript II reverse transcriptase (Thermo-Fisher Scientific). Real-time PCR were performed with primers listed in Supplementary Table 2, following a protocol previously described [27]. PCR primers were designed using Primer3web (https://primer3.ut.ee/). For each primer pair, amplification efficiency was measured using serial dilution of cDNAs prepared from cDNA mix of control and EX-527 treated samples, as described [35].

### EX-527 treatment *in vivo* in AT83 mice and in nude mice carrying AT29 cell grafts

AT83 mice and nude mice carrying AT29 cell grafts received 200 μL of either EX-527 (in 0.5% hydroxypropyl methylcellulose, 10 mg/kg of body weight) or its vehicle (0.5% hydroxypropyl methylcellulose) three times a week for 4 weeks by oral gavage, based on pharmacokinetic data previously obtained in mice [36]. Oral gavage with 10 mg/kg BW leads to maximal plasma concentrations of 10.5 ± 3.6 μM of EX-527 after less than one hour of gavage and to an average plasma concentration around 1.5 μM over 24 hours, with doubled concentrations in the brain at both times [36].

Swiss nude female mice (Charles River Laboratories, L’arbresle, France) were subcutaneously injected on the left flank with 2×10^6^ AT29 cells previously grown in complete medium and mixed with growth-factor enriched Matrigel® (Corning® #354248) (n=24 mice). The treatment by EX-527 or its vehicle was started once tumor volume reached at least 130 mm^3^ (n=12 mice/group). Graft size was assessed weekly with an electronic caliper, and volume was calculated according to the formulae, *length x width^2^/2*. For AT83 mice (CTR mice, n=4 and EX-527-treated mice, n=5), tumor size was followed once a week by three-dimensional ultrasound imaging using the Vevo LAZR high-frequency ultrasonic-photoacoustic imager (Vevo 2100, FUJIFILM VisualSonics) (Plateforme d’imagerie du vivant, Institut Cochin, Paris). Mice were placed on a support suitable for photoacoustic imaging which includes a mask for isoflurane inhalation, a heating plate equipped with a real-time cardiac, respiratory and temperature monitoring system. Flanks of the mice were shaved using commercial hair removal cream prior to applying the ultrasound gel for photoacoustic coupling interface between the ultrasound probe and the animal. Tumors were scanned in three dimensions (3D) with the help of an automated motor system. Tumor volume was established using VEVO LAB 3.1.1 software. Calculation of the tumor volume gain or loss was obtained by subtracting the measured volume at each week of treatment to the one measured at the beginning of the treatment. All experiments were performed in accordance with standard ethics guidelines and approved by Institutional Animal care and Use committee of the University Paris Cité and by the French Ministry of Agriculture (agreement #02193.02).

### Statistical analyses

Data was analyzed by Prism 6 (version 6.0, GraphPad Software). We used non-parametric tests when samples were not normally distributed as determined by Shapiro-Wilk normality tests. Depending on the experimental setting, we used Student *t-*test, parametric and non-parametric one-way ANOVA (Mann-Whitney, Kruskal-Wallis or Friedman) and two-way ANOVA. Data are shown as means ± SD or SEM and in scatter plot graphs with means. A *P*-value < 0.05 was considered as significant.

## Results

### SIRT1 is overexpressed in mouse and human GCT

We investigated whether SIRT1 expression could be altered during GC tumorigenesis by studying its subcellular localization between healthy ovaries and GCT harvested from patients. Our *in situ* analyses of human ovaries showed that SIRT1 was mainly expressed in oocytes, thecal cells and GC of growing follicles, with a predominant expression in GC nuclei (Figure 1Aa), as reported [37]. We observed a consistent expression of SIRT1 in primary and metastatic GCT (Figure 1Aa), with a predominant nuclear localization (insets in Figure 1Aa). We next investigated whether SIRT1 protein abundance could be altered in GCT, by comparing its relative expression levels between healthy luteinized GC and primary GCT at an advanced stage by Western Blot [3] (Figure 1Ab). Our results showed that SIRT1 abundance increased by ∼2 fold in GCT as compared with that in luteinized GC.

**Figure 1:**
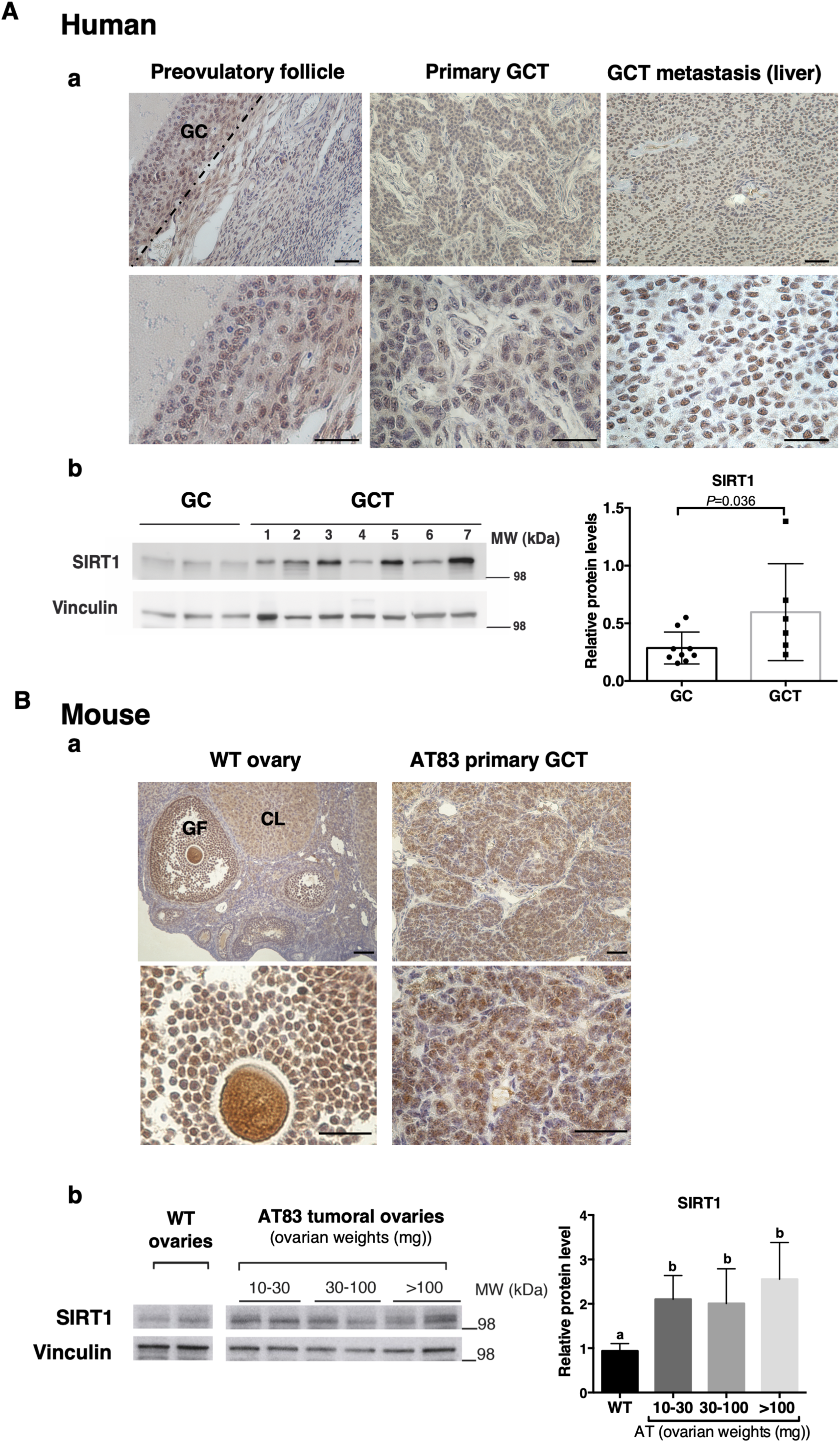
SIRT1 is overexpressed in tumoral granulosa cells. **(A)** Comparison of SIRT1 expression levels between healthy and tumoral GC in women. (a) SIRT1 immunodetection in healthy and tumoral or metastatic GC (dark yellow staining; hematoxylin counterstaining in blue; magnification in the inset). On the left panel is shown SIRT1 immunodetection in the ovary of a 32-year-old patient, with a prominent nuclear staining in GC of a preovulatory follicle. In the middle and right panels, primary GCT and GCT metastasis in the liver, respectively show a consistent SIRT1 nuclear expression. Scale bars: 50 μm. (b) Determination of SIRT1 and vinculin (used as a loading control) expression levels in GC and advanced primary GCT by Western blotting. The graph on the right shows the relative SIRT1 protein levels obtained after quantification of band intensities of SIRT1 and vinculin by ImageJ and normalization to vinculin for each sample. The number of samples for each group is the following: GC, n=9; GCT, n=6. Bars represent the means ± SD of normalized data. Data were analyzed by a non-parametric Mann-Whitney test. **(B)** Comparison of SIRT1 expression levels between healthy and tumoral GC in mice. (a) Immunodetection of SIRT1 in normal and tumoral mouse ovaries. In the wild-type mouse ovary counterstained by hematoxylin (blue), SIRT1 staining (dark yellow) is visible in GC and oocytes of growing follicles (GF), and in luteal cells of corpora lutea (CL). There is a prominent nuclear staining in GC (inset). In the primary GCT from an AT83 mouse, SIRT1 is observed in tumoral GC and displays a nuclear localization. Scale bars: 50 μm. (b) Representative Western blot showing SIRT1 and vinculin (used as a loading control) abundance at different stages of tumor growth in the AT83 mouse. The graph on the right shows the relative SIRT1 levels obtained after quantification of band intensities of SIRT1 and vinculin by ImageJ and normalization to vinculin for each sample. The number of samples for each group are the following: WT ovaries, n= 7; AT83 tumoral ovaries: 10-30 mg, n= 6; 30-100 mg, n=5; > 100 mg, n=5. Bars represent the means ± SD of data. Data were analyzed with non-parametric one-way ANOVA test (Kruskal-Wallis), with different letters indicating significant differences between groups (*P*<0.05).

To get more insights into the evolution of SIRT1 abundance during GC tumorigenesis, we took advantage of a transgenic mouse model of GCT that recapitulates major pathological features observed in patients, i.e., the AT83 mouse [27]. In this mouse model, we have defined three stages of tumor growth related to morphological criteria and ovarian weights: 10-30 mg (initial stage: newly developed/small GCT co-existing with corpora lutea and growing follicles), 30-100 mg (intermediate stage: small to medium GCT, few growing follicles and corpora lutea), and above 100 mg (late stage: large GCT) [27]. As in humans, SIRT1 was immunodetected in healthy GC from wild-type (WT) ovaries and in GCT of AT83 mice (Figure 1Ba). Importantly, SIRT1 protein abundance increased approximately 2-fold in AT83 ovaries of all categories compared to WT ovaries (Figure 1Bb). Furthermore, there was no significant change in SIRT1 abundance between the initial and final stages of tumor growth. These observations suggest that SIRT1 is overexpressed in tumoral GC *versus* GC from the earliest stage of tumorigenesis.

### Pharmacological inhibition of SIRT1 by EX-527 reduces cell growth in three GCT cell lines by affecting cell survival or cell proliferation

We next assessed *in vitro* the effect of pharmacological SIRT1 inhibition on common biological events related to tumorigenesis, i.e., cell growth, proliferation, viability, apoptosis and migration capacity, in different GCT cell lines. We used KGN (human, metastatic tumor), COV434 (human, primary tumor), and AT29 (mouse, primary tumor) cells, which display distinct molecular alterations found in GCT, such as *FOXL2 C402G*, *TERT* promoter and *KMT2D* mutations (KGN cells) [10,11], FOXL2 down-regulation (COV434 cells) [38], and inactivated P53/Rb pathways (AT29 cells) [27]. These three GCT cell lines expressed SIRT1 protein, as shown by Western blot (Figure 2A). The treatment by EX-527 at either 20 or 50 μM, chosen based on previous studies [18,21], significantly inhibited the activity of recombinant SIRT1 protein *in vitro* by ∼ 95%, with a dose-dependent effect (Figure 2B). SIRT1 activity was efficiently inhibited by ∼80-95% at 20 and 50 μM, with a dose-dependent effect in COV434 cells (Figure 2B). We therefore selected the highest dose of 50 μM for subsequent *in vitro* studies.

**Figure 2:**
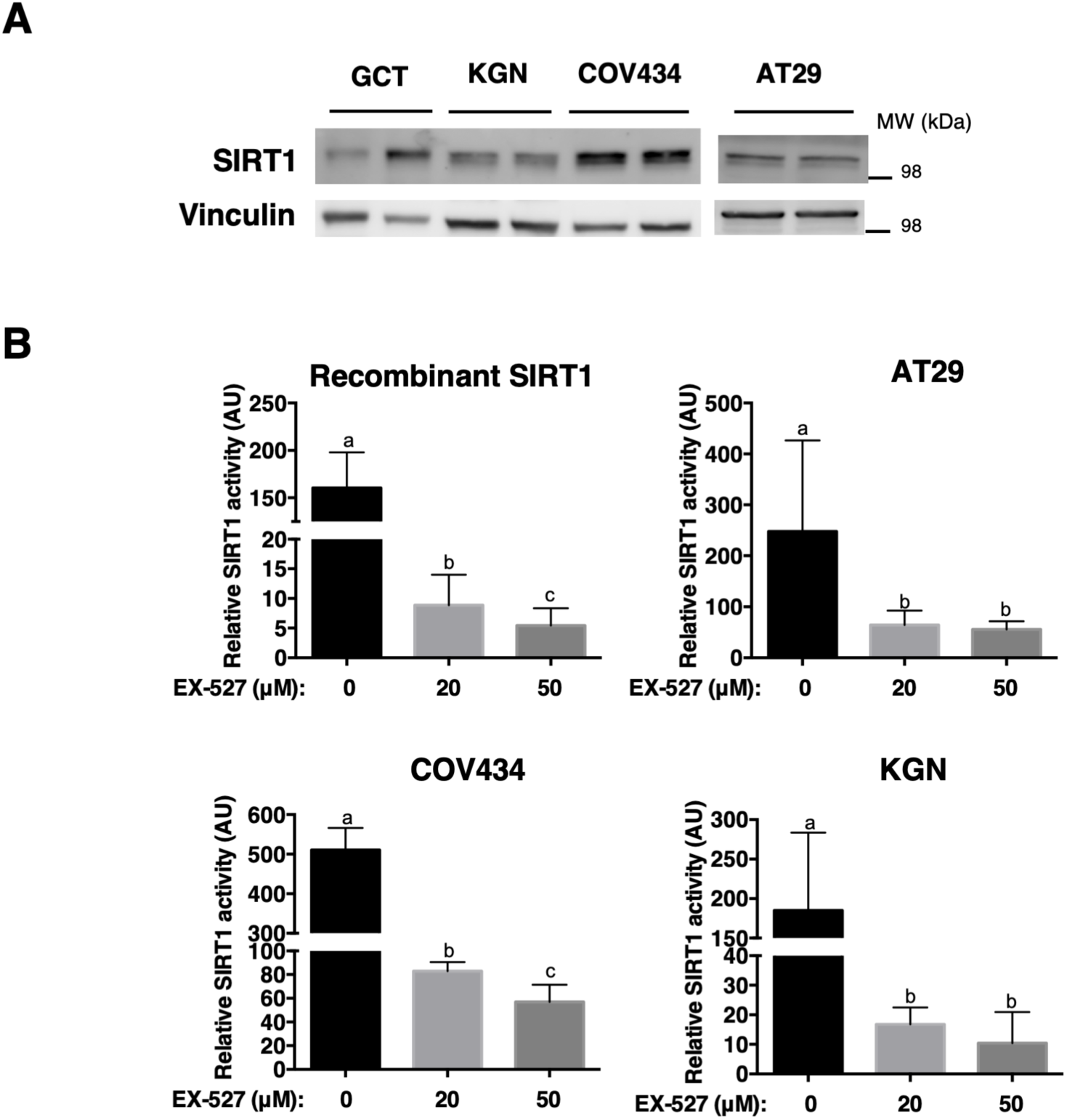
SIRT1 is expressed in different GCT cell lines and its activity is efficiently inhibited by EX-527 *in vitro.* **(A)** Determination of SIRT1 expression in human GCT and in three GCT cell lines of human (KGN, COV434 cells) and mouse (AT29 cells) origins, by Western blot assays. SIRT1 is expressed in the three cell lines. Vinculin is used as a loading control. **(B)** Effect of EX-527 at 20 and 50 μM on the activity of recombinant SIRT1 and endogenous SIRT1 in AT29, KGN and COV434 cells, as measured with a specific SIRT1 activity assay. The two used concentrations efficiently inhibit recombinant and endogenous SIRT1 activity in the three cell lines, with a dose-dependent inhibition observed for recombinant SIRT1 activity and COV434 cells.

We next evaluated the impact of EX-527 treatment on the growth of GCT cells by MTT assays (Figure 3A). As shown in Figure 3A, cell growth was significantly inhibited in the three cell lines, starting 24 hours after treatment in KGN cells and after 48 hours in COV434 and AT29 cells. As cell growth is the result of cell proliferation and survival, we then determined which of these processes was affected by the treatment. The fraction of viable cells decreased significantly upon EX-527 treatment in AT29 cells (by ∼50%), but not in KGN or COV434 cells (Figure 3B). Consistent with these data, the analysis of caspase-3/7 activity showed that the treatment increased apoptosis by ∼20% in AT29 cells but not in the two other cell lines (Figure 3C). Studies of BrdU incorporation indicated that cell proliferation did not significantly change in AT29 cells in response to EX-527, whereas it decreased significantly by 12% and 20% in KGN and COV434 cells, respectively, compared to the control condition (Figure 3D).

**Figure 3:**
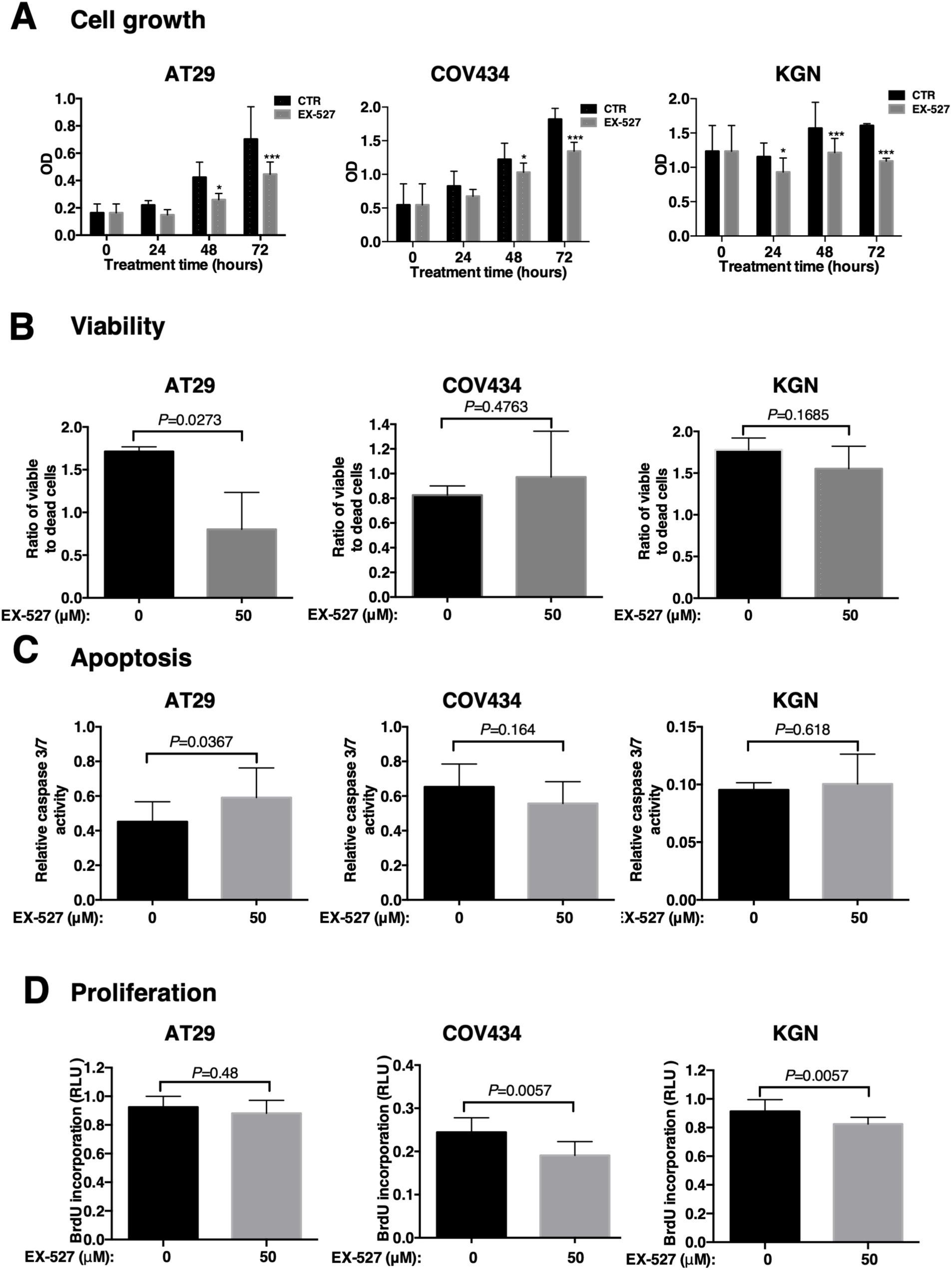
EX-527 induced inhibition of SIRT1 activity results in decreased cell growth by limiting cell viability and cell proliferation, and by increasing cell apoptosis. **(A)** Cell growth in response to EX-527 treatment at 50 μM for 24, 48 and 72 hours compared to control vehicle (CTR) by MTT assays in AT29, COV434 and KGN cells. Optical density (OD) measurement correlates with cell number. Graphs show the mean results obtained from independent experiments (n=3 for KGN and COV434; n=6 for AT29 cells). **(B)** Cell viability in response to a 24-hour treatment by EX-527 at 50 μM in AT29, COV434 and KGN cells. It was determined by using fluorescent substrates reflecting the quantity of live and dead cells. Graphs show the mean results obtained from independent experiments (n=3 for KGN and COV434; n=6 for AT29 cells). **(C)** Cell apoptosis was analyzed by measuring caspase-3/7 activity in the three cell lines treated with control vehicle (CTR) or EX-527 (50 μM) for 24 hours. Caspase-3/7 activity is represented as a relative luminescence unit (RLU), after normalization by the ratio of living to dead cell numbers for each sample. The number of experiments for each group are 3 for KGN and COV434 and 5 for AT29 cells. **(D)** Cell proliferation was determined by measuring BrdU incorporation by ELISA assays in AT29, COV434 and KGN cells treated with either control vehicle (CTR) or EX-527 (50 μM) for 24 hours. Optical density (OD) measurement correlates with BrdU incorporation levels. Shown are the results obtained from several independent experiments (KGN: n=9, COV434: n=7, and AT29 cells: n=4). In the graphs, the horizontal and error bars represent the means ± SD, respectively. Data in A were analyzed with paired two-way ANOVA. Data in B, C and D were analyzed by non-parametric Student t test (Kruskal-Wallis), with * *P* < 0.05, *** *P* < 0.001.

We also analyzed the effect of the SIRT1 inhibitor on cell migration, by performing transwell assays. We could not analyze this process in COV434 cells, as these cells do not have the ability to migrate [13]. As shown in Supplementary Figure 1, the treatment tended to decrease migration capacity in both cell lines, but this effect was not significant (*P*=0.125). Altogether, these data suggest that EX-527 treatment essentially affects GCT cell growth, by reducing either cell viability or cell proliferation.

### Transcriptome analyses revealed EX-527-induced alterations in oncogenic pathways in common in the three GCT cell lines despite their molecular heterogeneity

To get insights into the molecular pathways induced by EX-527 treatment, we analyzed the transcriptomes of the three cell lines by performing RNA-sequencing after a 24-hour exposure (Supplementary Table 1). PCA analyses revealed that the treatment affected gene expression in the three cell lines, with some variability between samples of the same treatment group (Supplementary Figure 2). By selecting a fold change (FC) at 2, EdgeR analyses identified more than 200 annotated differentially expressed ENSEMBL genes (DEGs) in each cell line (Table 2). The global transcriptional change between control and EX-527 treated cells is represented by a Volcano Plot for each cell line (Figure 4A). Comprehensive lists of differentially expressed genes in each cell line are given in Supplementary Table 3. We confirmed these data by performing RT-qPCR on a few DEG linked either to cell death for AT29 cells (*Parp14*, *Xaf1*) or to cell proliferation in KGN and COV434 cells (*CDCA3*, *FOS*, *GREM1*, *IL6*) (Figure 4B). We found no DEG in common between the three cell lines, even with FC>1.25, as shown in Venn diagrams (Figure 4C). However, there were a few overlapping genes positively or negatively regulated by EX-527 between AT29 cells and either KGN or COV434 cells, and between KGN and COV434 cells (Figure 4C).

**Figure 4:**
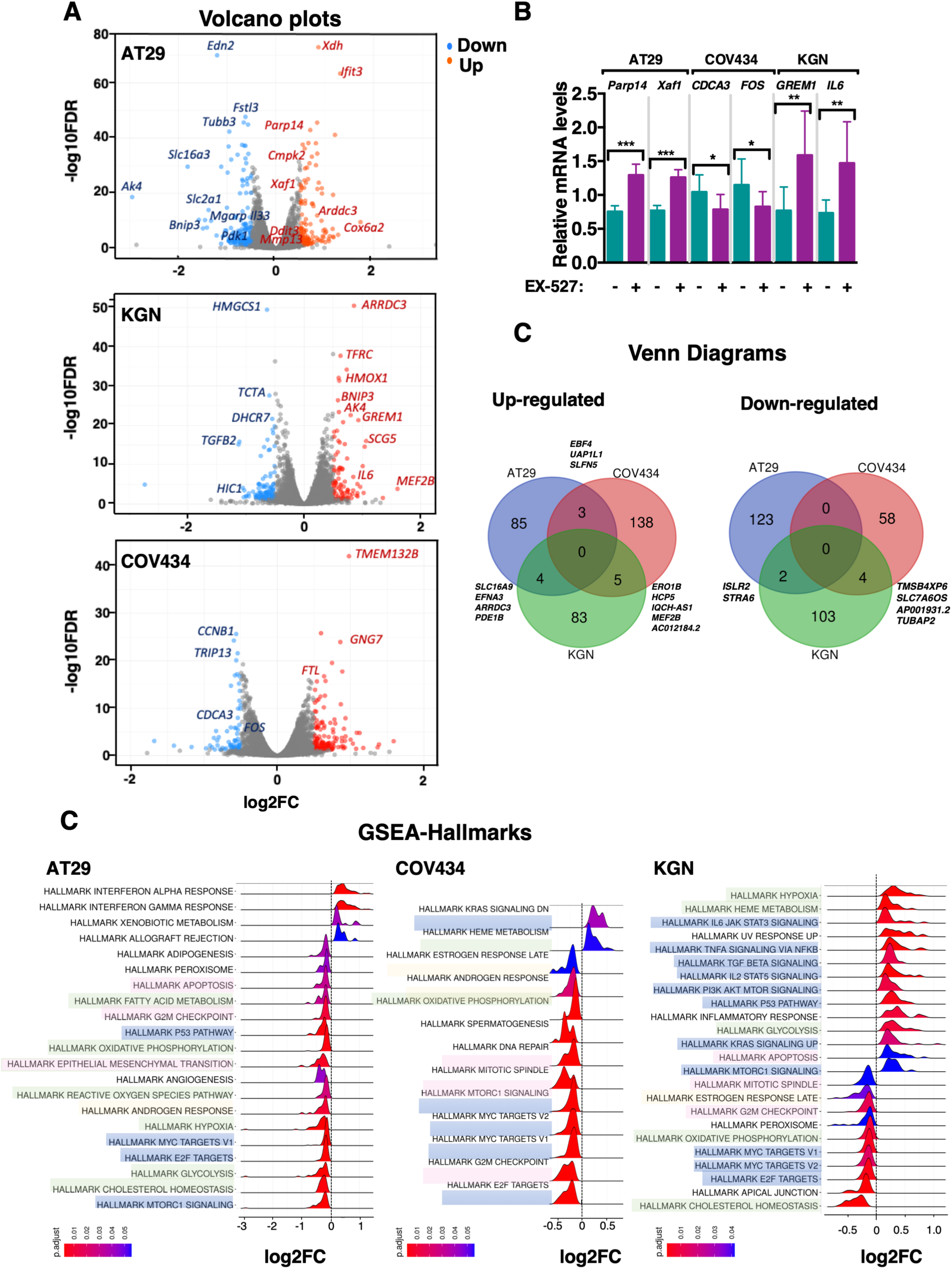
Transcriptome analyses by RNA-sequencing reveal EX-527-induced alterations of pathways in common in AT29, COV434 and KGN cells. **(A)** Volcano plots of DEG in cells treated by EX-527 treatment (50 μM) for 24 hours compared to control cells. A few up-(red dots) and down-regulated (blue dots) genes are indicated on each plot for AT29, COV434 and KGN cells. **(B)** Relative expression levels of selected genes found to be differentially expressed by RNA-sequencing after EX-527 treatment in each cell line. It was determined by quantitative real-time RT-PCR and normalized to the mRNA levels of *HPRT* in 9 RNA samples per group and per cell line, prepared in 3 independent experiments. **(C)** Venn diagrams showing the number and names of up- and down-regulated genes that overlap between cell lines. **(D)** Hallmarks found altered by Gene Set Expression Analysis (GSEA) in AT29, COV434 and KGN cells. The ridgeplots represent density plots of the frequency of log2FC values per gene within each set from significantly up and down-regulated pathways, for which *P*-adjust value was <0.05. The altered signaling pathways are highlighted in blue, altered cellular processes in pink, and metabolic pathways in green color.

**Table 2:** Total number of genes expressed and differential gene expression in AT29, COV434 and KGN cells.

| GCT cell line | Total number of genes<br>expressed | Differentially expressed genes |  |
| --- | --- | --- | --- |
|  |  | Up | Down |
| **AT29** | 14 161 | 132 | 192 |
| **COV434** | 14 110 | 143 | 66 |
| **KGN** | 13 372 | 92 | 109 |

To identify the pathways that were altered by EX-527 treatment, we used the Gene Set Enrichment Analysis (GSEA) and data were analyzed through the Hallmark Gene Sets database (Figure 4D). Drug treatment altered several oncogenic pathways in common among the three cell lines, i.e., E2F, Myc mTOR, which were all downregulated (highlighted in blue in Figure 4D). EX-527 treatment also altered the expression of genes related to the P53 pathway in AT29 cells (decrease) and in KGN cells (increase). Consistent with the observed effect of EX-527 on cell growth in the three cell lines, there was an alteration in G2/M checkpoint-related genes, and in those regulating apoptosis (AT29 cells) or the mitotic spindle (KGN and COV434 cells). There was also a significant alteration in genes linked to androgen and/or estrogen response, suggesting that the treatment could interfere with sex steroid signaling (Figure 4D).

### EX-527 treatment alters the expression of genes related to mitochondrial function and it increases reactive oxygen species in vitro

Interestingly, GSEA analysis of hallmarks and GO-biological processes revealed alterations in metabolic-related pathways including a decrease in oxidative phosphorylation pathway in the three cell lines, consistent with the previously reported repressing effect of EX-527 on ATP production in KGN cells [18] (Figure 4A and Figure 5A, Supplementary Figure 3). Specifically, EX-527 treatment decreased the expression of the mitochondrial ATP synthase subunits *ATP5* (shown as black stars in the heatmaps) in the three cell lines. It also affected the expression of *NDUF* genes, which encode the subunits of the enzyme NADH dehydrogenase (ubiquinone), part of the Complex I of the electron transport chain located in the mitochondrial inner membrane and the largest of the five complexes of the electron transport chain (shown as purple stars in the heatmaps) [39]. GSEA analysis also revealed that EX-527 down-regulated the expression of different subunits of cytochrome c oxidase of complex IV of the mitochondrial respiratory chain (*COX4*, *COX5*, *COX6*, *COX7*, *COX8*) and that of the cytochrome oxidase chaperone *COX17* that contributes to cytochrome c oxidase activity (shown as green stars in the heatmaps). None of the affected genes were of mitochondrial genome origin, despite their action on mitochondrial function. To get more insights into the impact of EX-527 treatment on the mitochondria, we performed additional analyses of DEG encoding proteins with strong support of mitochondrial localization using the MitoCarta3.0 database. There was an altered expression (FC>2) for 8 of them in AT29 cells but for only 1 of them in KGN cells and none in COV434 (Supplementary Table 4). However, when we analyzed our RNA-seq data with a FC set up at 1.25 instead of 2, we identified more than 10 DEG related to mitochondria function in each cell line (Supplementary Table 4). Analyses of genes (FC>1.25) regulated by Nuclear factor erythroid 2-related factor 2 (NRF2), a transcription factor that regulates cellular defense against oxidative insults through the expression of genes involved in oxidative stress response [40], revealed that a subset of them were differentially expressed between control and EX-527 treated cells (Supplementary Table 5). We also sought for a possible alteration in ferroptosis pathway, a type of cell death resulting in a decrease in antioxidant capacity and accumulation of lipid ROS in cells, ultimately leading to oxidative cell death [41]. We thus analyzed our RNA-seq data using the FerrDb website, a database that dedicates to ferroptosis regulators and ferroptosis-disease associations(http://www.zhounan.org/ferrdb/current/operations/enrichanalysis.html). However, only a small subset of genes exhibited altered expression in AT29 cells and KGN cells, and none in COV434 cells, even with FC>1.25 (Supplementary Table 5). We confirmed that the treatment significantly altered the expression of selected genes in the three cell lines, including those with FC comprised between 1.25 and 2, by performing RT-qPCR (Figure 5B).

**Figure 5:**
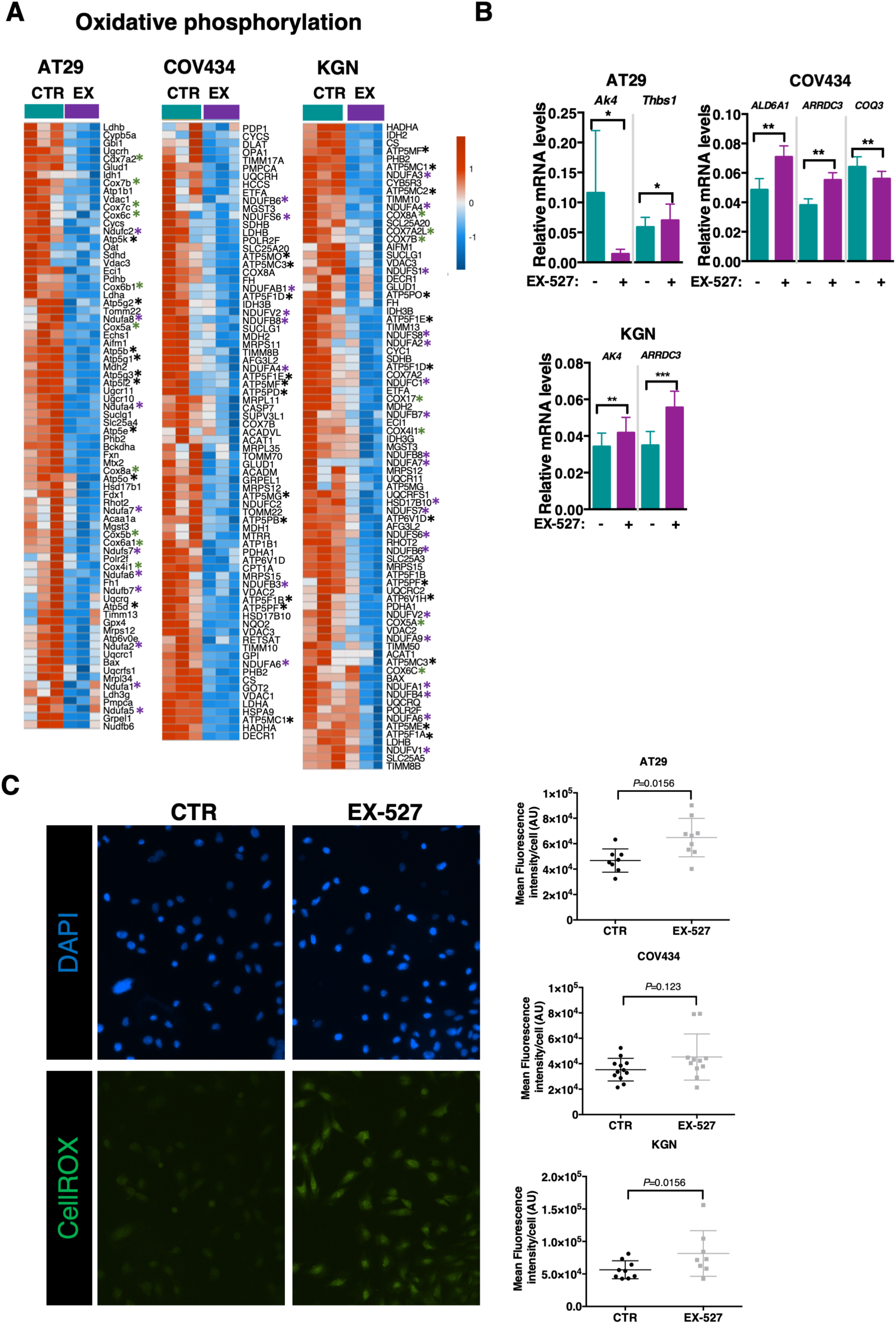
EX-527 treatment induces alterations in metabolic pathways in the three cell lines *in vitro*. **(A)** Heatmaps of GSEA showing that EX-527 treatment for 24 hours induces an overall decrease in genes belonging to gene sets related to oxidative phosphorylation. **(B)** Relative expression levels of selected genes related to mitochondrial pathway found to be differentially expressed after EX-527 treatment in each cell line. It was determined by quantitative real-time RT-PCR and normalized to the mRNA levels of *HPRT* in 9 RNA samples per group and per cell line, prepared in 3 independent experiments. **(C)** Effect of EX-527 treatment on ROS generation. Shown is the green fluorogenic signal produced by ROS in AT29 cells stained by DAPI (blue) to visualize cell nuclei. A higher green fluorescence intensity is observed in the nucleus of AT29 cells after EX-527 treatment than in control (CTR) conditions. Graphs show the quantification of the mean fluorescent intensity determined by ImageJ in the three cell lines from at least three independent experiments. The *P* values obtained by non-parametric Student t-test are shown in the graphs.

The alterations induced by EX-527, and notably those on mitochondrial function, led us to investigate whether the treatment had an effect on reactive oxygen species (ROS). We measured ROS *in vitro* using fluorogenic probes being weakly fluorescent in a reduced state and exhibiting strong fluorogenic signal upon oxidation (illustrated for AT29 cells in Figure 5C). There was a ∼1.3-fold increase in fluorescence intensity in the three cell lines after EX-527 treatment as compared with the control condition, which was significant in AT29 and KGN cells (*P*=0.0156) but not in COV434 cells (Figure 5C).

### Pharmacological inhibition of SIRT1 reduces GCT growth in two mouse models of the disease by preventing cell survival and proliferation

We then tested *in vivo* the effect of the inhibition of SIRT1 activity by EX-527 in immunodeficient mice carrying subcutaneous allografts of AT29 cells on the left flank (Figure 6Aa, c). We started treatment in mice harboring a graft volume > 130 mm^3^, reached about 10-11 weeks after cell injection. EX-527 did not alter body weight throughout the experiment duration, indicating that it had no major adverse effects, consistent with the lack of reported toxicity of this drug [21] (Figure 6Ab). In vehicle-treated mice, graft volume increased after one week (range: 163 to 450 mm^3^) and it remained stable over the next three weeks (Figure 6Ad). In contrast, in EX-527-treated mice there was a rapid reduction in tumor volume that became significant after 4 weeks of treatment (Figure 6Ad). Accordingly, at dissection graft weights were significantly lower (by ∼25%) in mice that received EX-527 as compared with those treated with the vehicle (Figure 6Ae).

To gain more insights into the efficacy of EX-527 in repressing GCT growth *in vivo*, we used AT83 mice developing endogenous GCT. We selected mice of our colony aged >12 months, as 100% of them display GCT at this age. Because GCT growth is asynchronous between AT females of the same age and the size of GCT developing on the ovaries of the same mouse can greatly vary [27], we monitored tumor growth from the onset of treatment by *in vivo* imaging using a 3-D ultrasound system with the VEVO LAZR technology (Figure 6Bb). Similar as in nude mice, EX-527 treatment did not impair the overall health of the mouse, as shown by the body weight which remained stable over time and comparable between the two groups (Figure 6Bc). At the onset of the experiment, tumor volume greatly differed between mice and between the two ovaries of the same mouse, as expected [27] (Supplementary Table 6). In vehicle-treated AT83 mice, there was a progressive gain in tumor volume in all analyzed tumoral ovaries over the four-week follow-up (Figure 6Bd). This was markedly different in EX-527-treated AT83 mice, in which only 3 out of 10 ovaries showed tumor growth, while 3 out of 10 did not show any growth and 4 out of 10 even showed a regression in tumor volume (Supplementary Table 6). Overall, EX-527 induced a significant decrease in GCT growth that became significant after 3 weeks of treatment (Figure 6Bd).

We then analyzed the effect of EX-527 treatment on cell apoptosis and proliferation using Western blot assays with protein lysates prepared from AT29 cell grafts collected after 4 weeks of EX-527 treatment. These grafts are homogeneous in cellular population, at the difference of GCT from AT83 mice. Analyses of the abundance of an initiator caspase contributing to extrinsic apoptosis, i.e., caspase-8, revealed that the ratio of its cleaved (active) to un-cleaved (inactive) forms significantly increased from approximately 2-fold in the grafts from EX-527-treated mice as compared with those in control mice, indicating that the treatment induced cell apoptosis (Figure 6C). In addition, the relative expression levels of a cell cycle-promoting gene, i.e., cyclin D2, decreased approximately 1.4-fold in the grafts of EX-527 treated mice when compared with those in controls (Figure 6C). These data suggest that EX-527 treatment led to the involution of AT29 cell graft by promoting cell apoptosis and by inhibiting cell proliferation.

**Figure 6:**
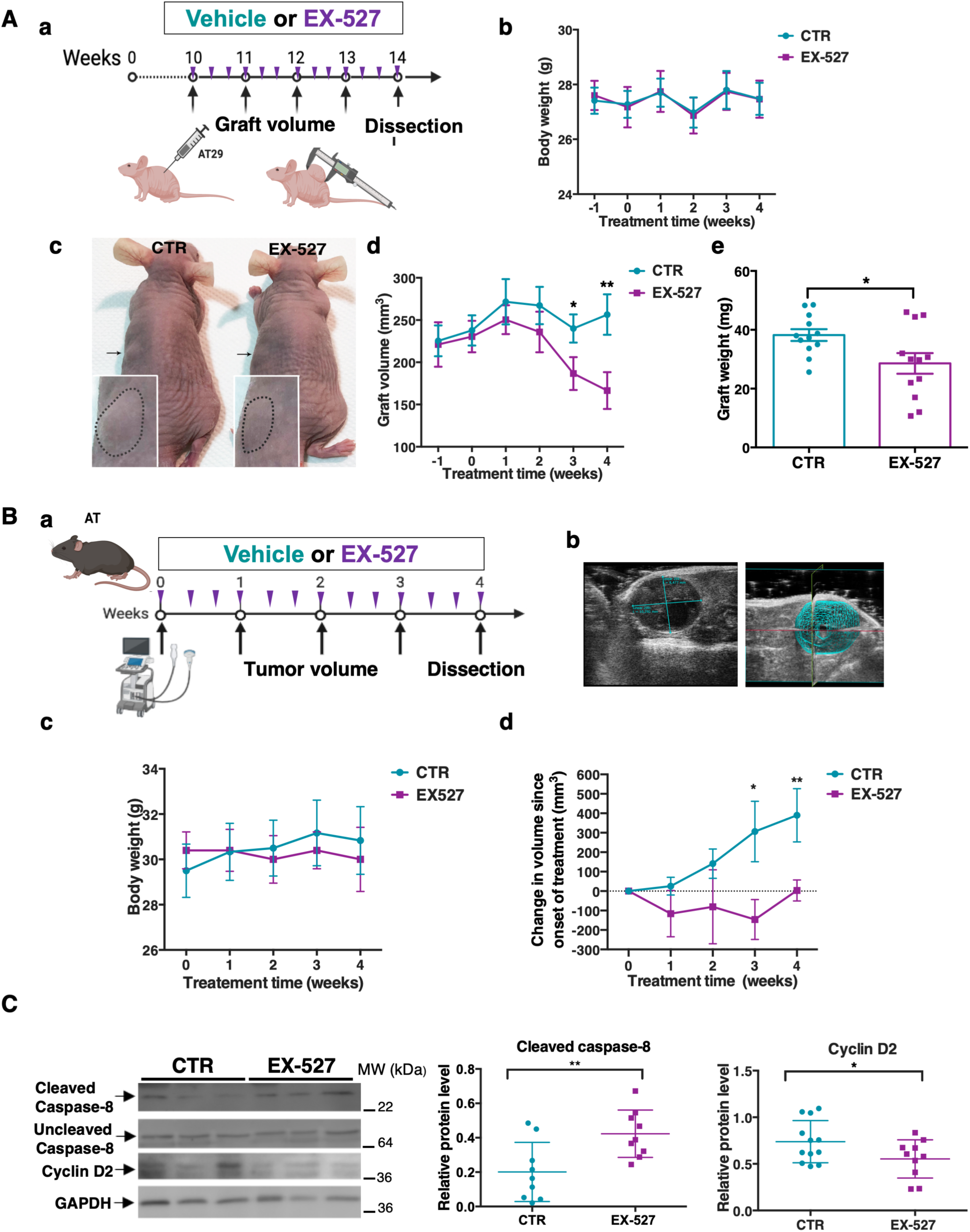
SIRT1 inhibition by EX-527 treatment decreases tumor growth in two mouse models of GCT. **(A)** Response to EX-527 on the growth of grafts prepared from AT29 cells in nude mice. (a) Schematic representation of the procedure used to monitor the growth of the graft. Purple triangles represent the day of oral gavage with either vehicle or EX-527, and black arrows show the days of measurement of graft size with calipers. (b) Body weights of AT29 cell-grafted nude mice 1 week before and over the 4 weeks of EX-527 treatment. The number of mice for each group are n=12. Dots and error bars represent the means ± SEM, respectively. (c) Representative photographs of the subcutaneous grafts in immunodeficient mice after 4 weeks of treatment with EX-527 or its control (CTR) vehicle. Black arrows show grafts at the end of treatment. An enlargement of the grafts is shown to visualize the size of the tumor. (d) Graph shows the mean of the volumes of the grafts 1 week before and over the 4 weeks of EX-527 treatment. The number of samples for each group is n=12. Dots and error bars represent the means ± SEM, respectively. (e) The graph shows the mean graft weights at the end of the 4-week treatment with EX-527 or its control vehicle. Individual values are shown in scatter dot plots with means of normalized data (horizontal bars) with n=12 grafts for each group. *, *P* < 0.05. Data in b and d were analyzed with two-way ANOVA test with * *P* < 0.05 and ** *P* < 0.01. Data in e were analyzed by Student t-test. **(B)** Response to EX-527 of GCT growth in AT83 transgenic mice. (a) Schematic representation of the procedure used to investigate the response of GCT growth to EX-527 or its vehicle in AT83 mice. Purple triangles represent the time of oral gavage with EX-527 or its vehicle, and black arrows show the time of three-dimensional tumor volume recording by ultrasound imaging. (b) Bi-dimensional (left) and three-dimensional (right) views of GCT from AT83 mice acquired by the VEVO ultrasound system. (c) Body weights over the 4 weeks of treatment with EX-527, or its control vehicle. The number of mice is n=4 for CTR and n=5 for EX-527. (d) Weekly evolution of the gain in tumor volume since the start of treatment. The calculation of the change in volume was obtained by subtracting the volume measured at each week of treatment from that at the start of treatment. The number of GCT is n=7 in CTR, and n= 10 in EX-527-treated group. Data in b and d were analyzed with two-way ANOVA test with * *P* < 0.05 and ** *P* < 0.01. **(C)** Determination of the levels of cleaved and uncleaved forms of caspase-8 and cyclin D2 in AT29 cell grafts of mice treated with EX-527 or their control vehicle (CTR), by Western blot assays. Cleaved caspase-8 levels were normalized to those of uncleaved caspase-8, whereas cyclin D2 levels were normalized to those of GAPDH for each sample. Representative immunoblots are shown. The number of samples for each group are n=9 for CTR and n=8-9 for EX-527 and individual values are shown in scatter dot plots with means of normalized data (horizontal bars). Band intensities were quantified using ImageJ. For all these data, the horizontal and error bars represent the means ± SD, respectively. Data were analyzed by Student *t*-test with * *P* < 0.05, ** *P* < 0.01.

## Discussion

SIRT1 acts as an oncogene in several types of cancer where the benefit of its inhibition is being evaluated [14], but its role in the pathogenesis of GCT remained unclear. Here, by performing *in vitro* studies on various cell lines and *in vivo* studies in the mouse model together with translational studies on human samples, we present evidence that SIRT1 is overexpressed early in GC tumorigenesis and that repressing its activity could provide therapeutic benefits in patients with GCT.

Our comparison of SIRT1 abundance revealed its approximatively two-fold higher expression in GCT compared to luteinized GC collected from patients, with overexpression found in 100% of cases. The use of AT83 mice, in which the abundance of SIRT1 can be linked to GCT growth stage, showed that SIRT1 expression increases markedly at the very beginning of tumor growth, when the proportion of the tumor is still low in the ovary. Although we were not able to have perfect controls for human and mouse studies (the human GC used are luteinized following the stimulation protocol with a possibly induced change in SIRT1 abundance [42], and the WT ovaries contain several cell types including GC), our results suggest that SIRT1 becomes abnormally elevated early in GC tumorigenesis and remains elevated throughout tumorigenesis, thus possibly contributing to the oncogenic process. Unlike the present study, a previous report showed that SIRT1 is expressed in only 70% of GCTs [18]. This apparent discrepancy may come from the different number of samples (7 in our study *versus* 72 in [18]) and/or methodological approaches (Western blot here *versus* immunohistochemistry). This may also come from the different stages of the GCT analyzed, which were all at an advanced stage in our study, unlike that in [18]. High SIRT1 expression is found in some cancers, including prostate and breast cancers and could be of bad prognosis [17,43]. However, the possible correlation between SIRT1 expression and prognosis is still unclear in patients with GCT, given the long history of the disease [18].

Importantly, our in-depth work provides evidence that repressing SIRT1 activity in GCT cells could efficiently reduce tumor burden. EX-527 treatment reduced the growth of three GCT cell lines *in vitro,* regardless of their type of molecular alterations, although by affecting distinct cellular processes. Indeed, this action of EX-527 mainly resulted from reduced cell viability by apoptosis induction in AT29 cells, whereas it preferentially resulted from decreased cell proliferation in KGN and COV434 cell lines. The effect of EX-527 on KGN cell proliferation has been suggested previously, and may result from a decreased capacity of cells to exit or to enter S-phase although additional studies would be required to firmly establish it [15,18]. Although EX-527 treatment did not change the expression of the same genes in all three cell lines, this compound similarly impacted pathways related to the regulation of DNA repair, apoptosis, cell cycle (G2/M checkpoint and mitotic spindle), and oxidative phosphorylation. In addition, the treatment affected the expression of targets of mTOR and of the transcription factors Myc and E2F, which are all known to regulate the expression of genes involved in cell cycle and cell apoptosis [44–46]. mTOR is a signaling hub, sensing and integrating environmental and intracellular nutrient cues to coordinate cell growth, proliferation and apoptosis depending on cell type. It is frequently overactivated in cancers [47]. Myc is a major driver of cancer, where it is commonly overexpressed. Its inhibition or removal *in vitro* and *in vivo* leads to growth arrest of cancer cells [46]. The transcriptional activity of E2F, which is inhibited by the retinoblastoma protein (pRb), is frequently enhanced in neoplastic cells, leading to the inability to exit the cell cycle after DNA damage, which can cause genomic instability, promote malignant progression, and reduce drug sensitivity [48]. Besides, high expression of the transcription program of E2F correlates with poor prognosis in various types of cancer [49,50]. Combined E2F activation and loss of p53 function in GC may lead to GCT development, highlighting the key contribution of these pathways in this disease [27]. It is, thus, tempting to speculate that the inhibition of mTOR, Myc and E2F transcription program by EX-527 treatment results in the observed reduction in cell growth *in vitro* in the three GCT cell lines. However, it is possible that inhibiting these pathways could also modulate mitochondrial function and cell metabolism, notably by impacting the expression of mitochondria-associated genes (for review: [44–46]). Our RNA-seq analyses suggest that EX-527 treatment regulated mitochondria-associated genes, and in particular those involved in oxidative phosphorylation, including ATP synthase, cytochrome c oxidase and NADH dehydrogenase, which are all components of the mitochondrial respiratory chain. This effect of EX-527 on oxidative phosphorylation has been previously reported in KGN cells, where it led to an 80% decrease in ATP production following a 24 hour-treatment by EX-527 at 50 □M [18]. Oxidative phosphorylation is key for maintaining sustained growth and immortality of tumors by promoting cell metabolism [39]. We, thus, hypothesize that this effect of the treatment contributes to decrease GCT cell growth by enhancing apoptosis or proliferation, depending on cell lines. Furthermore, our analyses of GO biological processes suggest that EX-527 affected various aspects of cell metabolism such as the biosynthesis, metabolism and transports of lipids, amino acids, nucleotides and carbohydrates. In addition to the alteration in cell metabolism, we postulate that the inhibition of SIRT1 activity by EX-527 treatment increased ROS levels in GCT cells, as seen in AT29 and KGN cells. Since cells typically undergo cell cycle arrest and enter the G_0_ phase or may undergo apoptosis upon high exposure to ROS [51], we hypothesize that the observed effect of EX-527 on GCT cell proliferation and survival could have resulted from increased oxidative stress. The increase in ROS levels following SIRT1 inhibition may be reflective of the impairment of the electron transport chain, as electrons leaks are the primary source of oxidative stress and resulting cellular damages. These results suggest that, although EX-527 treatment has a different impact on gene expression in the three cell lines, it ultimately affects the same pathways to reduce cell growth.

Noteworthy, this compound halted tumor progression *in vivo* in two mouse models of GCT. Since obtaining grafts of patient GCT or human GCT cell lines to nude mice is challenging due to low tumor-take rate and slow growth, we tested EX-527 effects on a new GCT mouse model carrying AT29 cell grafts and in a transgenic mouse model recapitulating major features of GCT [27]. The growth of AT29 cell grafts was rather slow in vehicle-treated mice, indicative of a low aggressiveness, as observed in most patients. Treatment by EX-527 three times per week reduced graft volume, an effect that became significant after 4 weeks. This beneficial effect of the treatment was also observed in AT83 mice during close monitoring of individual tumor volume by 3-D ultrasound imaging. It revealed that while tumor growth increased over time in vehicle-treated mice, it remained stable in most EX-527 treated mice. We currently do not know why the treatment was not effective in reducing the growth of all tumors, but we postulate that non-responding tumors may have poor vascularization. Our molecular studies in AT29 cell grafts showed that EX-527 treatment promoted cell death, as shown by increased ratios of cleaved to uncleaved caspase-8, and decreased cell proliferation as suggested by the decrease in the relative levels of Cyclin D2. These findings are in line with those obtained *in vitro* in GCT cell lines.

Taken together, our data suggest that SIRT1 may be aberrantly activated in GCTs to stimulate proliferation and suppress apoptosis by regulating a number of oncogenic pathways and cell metabolism. Inhibition of its activity by EX-527 treatment could effectively limit the growth of tumor and metastatic GC, whatever their molecular alterations. Given the great heterogeneity of molecular alterations beyond the C402G *FOXL2* mutation in GCT, the use of a selective and well-tolerated SIRT1 inhibitor with broad action on GCT cells may represent a relevant opportunity for the clinical management of patients.

## Acknowledgments

The authors wish to thank Dr Bruno Querat (Université Paris Cité, BFA unit, France) for providing SIRT1 antibody, Dr Anne Mayeur (Service de Médecine de la Reproduction et Préservation de la Fertilité, Hôpital Antoine Béclère, Clamart, France) for providing human GC. The authors acknowledge Dr N di Clemente (Centre de Recherche Saint-Antoine, Inserm U938, Paris) for providing AT29 cells and AT83 transgenic mice. RNA-seq was carried out by the ICM Data Analysis Core platform. They gratefully acknowledge Yannick Marie, Emeline Cherchame and Justine Guegan for their assistance on RNA-seq analyses. They would also like to thank the staff of the Buffon Animal Core Facility, and in particular Dr Isabelle LeParco (Université Paris Cité, France) and the “Plateforme Imagerie du Vivant” (Institut Cochin, France).

## Authors’ contributions

VC, EA, MMD, FP, SC, AP and CJG performed experiments. VC, EA, and CJG analyzed and interpreted the data. CPD, UW, HJ and CJG performed bioinformatic analysis of RNAseq data. AL provided biopsies from patients with GCT. CJG supervised the study and wrote the manuscript. VC and SC contributed to manuscript redaction. All authors read and approved the final manuscript.

## Ethics approval and consent to participate

All women were fully counseled, and informed consent was obtained in writing from all participants. This investigation received the approval of the Internal Institutional Review Boards from Antoine Béclère Hospital (granulosa cells) and Institut Gustave Roussy (GCT). The study was performed in accordance with the Declaration of Helsinki for Medical Research involving Human Subjects (2013 revision). The investigation received the approval of our internal institutional review board, IRB Blefco-IORG0010582, and is registered under number “2021-1”. Date of the first subject enrollment is January 2021.

All experiments in mice were performed in accordance with standard ethics guidelines and approved by Institutional Animal care and Use committee of the University Paris Cité and by the French Ministry of Agriculture (agreement #02193.02). A completed <u>ARRIVE2.0</u> checklist is provided as supplementary file.

## Consent for publication

N/A

## Data availability

RNAseq data are available in a public, open access repository, under accession number GSE252773.

