## Supplementary Figures for "Preclinical evaluation of pharmacological inhibition of SIRT1 on the growth of tumoral and metastatic granulosa cells"

**Supplementary figures & tables**

### Pharmacological inhibition of SIRT1 limits the growth of tumoral and metastatic cells from ovarian granulosa cell tumors by impacting several oncogenic pathways and cell metabolism

**Authors:** Victoria Cluzet, Eloïse Airaud, Marie M Devillers, Florence Petit, Alexandra Leary, Alice Pierre, Haojian Li, Chi-Ping Day, Urbain Weyemi, Stéphanie Chauvin, Céline J Guigon

**Supplementary Figure 1:** Capacity of migration of AT29 and KGN cells using the principle of Boyden chambers. Cells were seeded in transwells (fluoroblocks for AT29 cells, regular transwells for KGN cells) as described in *Supplementary Materials and Methods*. Data are the mean of three to five independent experiments (+/- SD) and were analyzed by non parametric Student *t*-test.

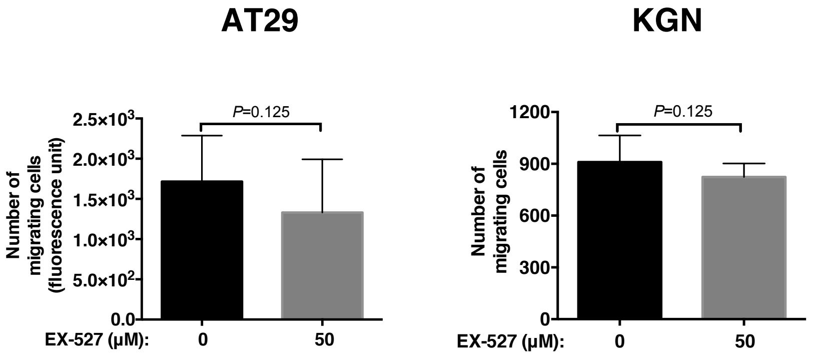

**Supplementary Figure 2**: PCA analysis of the samples run for RNA-seq. For AT29 and COV434 cells, light blue dots show control samples and dark blue dots those treated by EX-527. For KGN cells: light green dots show control samples and dark green dots those treated by EX-527

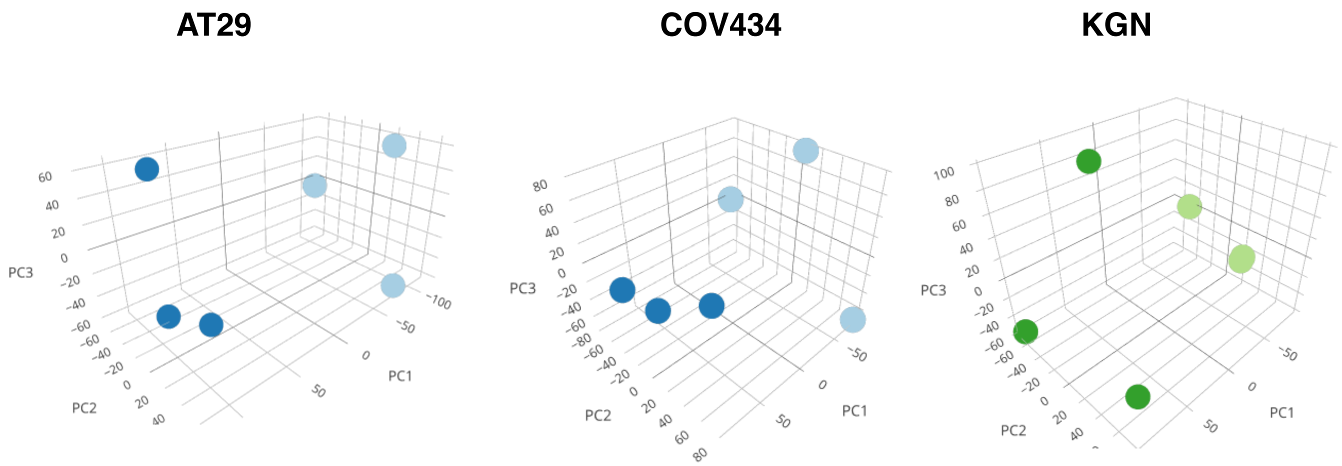

### Supplementary Figure 3:

**Gene ontology-biological processes**

The Dotplot depicts the normalized enrichment scores (NES) (enrichment score given normalized based on the number of genes in the gene set). The dots are colored by the adjusted *p*-value and their size is proportional with the size of the gene-set.

#### AT29 cells

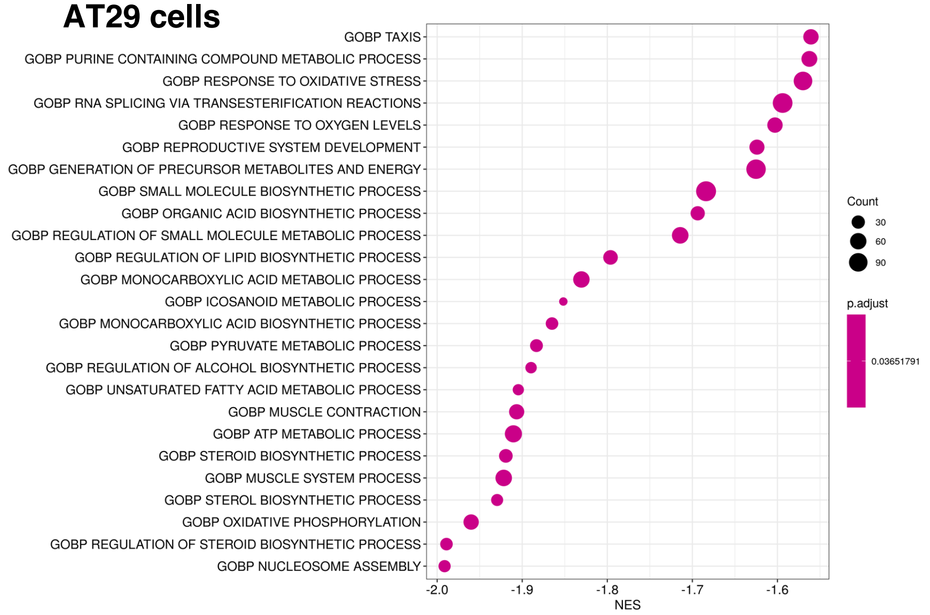


#### KGN cells

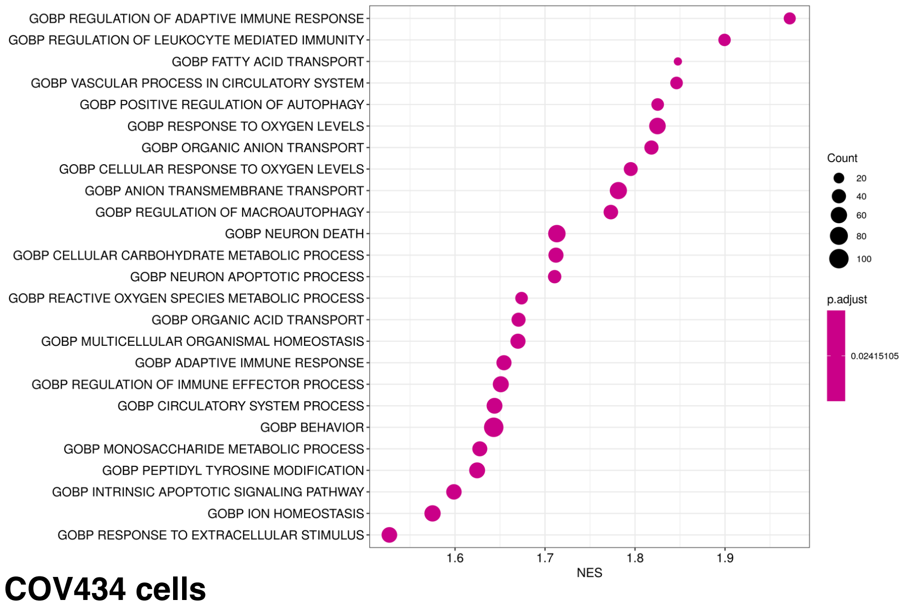


#### COV434 cells

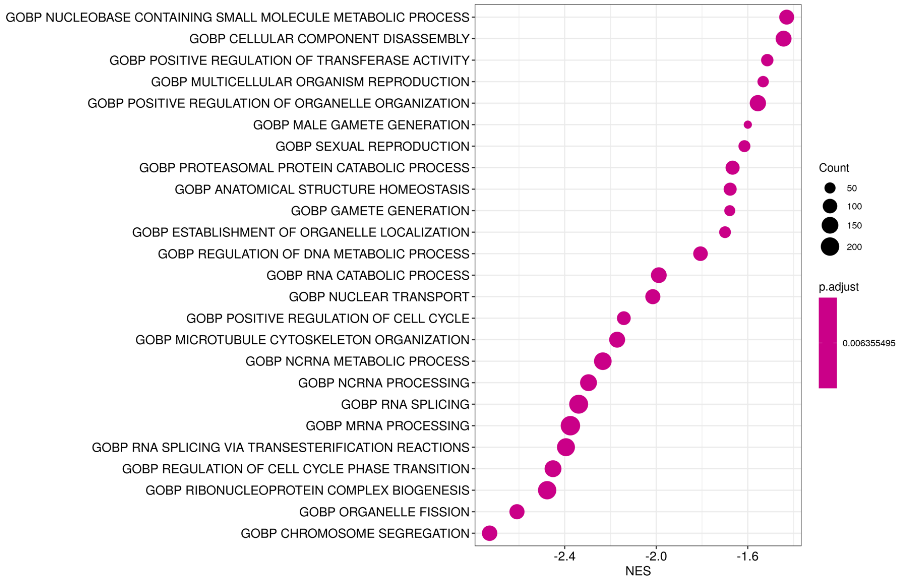

**Supplementary Table 1 (large Excel file of RNA-seq data):** global gene expression in human (KGN, COV434) and mouse (AT29) cell lines, markers of GC in human cell lines

**Supplementary Table 2:** list of primers used for qPCR on human (KGN, COV434) and mouse (AT29 cells) GCT cell lines

| | Forward | Reverse |
|---|---|---|
| **Human primers** | | |
| *AK4* | TGGATTCACCCTCCTAGCGGAA | CTGTCTTAGCCTGGCAGCAACT |
| *ARRDC3* | CTAGAAACGCCGGCTCCAAT | GTAGCGAGTGGTGTCTGTGG |
| *ALDH6A1* | AGCAGGTCTTGCTCCGCTATCA | GGATGTCACACTACAGGCATGC |
| *CCNB1* | AATAAGGCGAAGATCAACATGGC | TTTGTTACCAATGTCCCCAAGAG |
| *GREM1* | ATGTGACGGAGCGCAAATAC | TGGATATGCAACGACACTGC |
| *CDCA3* | ATTGCACGGACACCTATGA | TGTGGGCTGTCTTGCTTC |
| *COQ3* | AGTTTCAGGTACCCTTGGGC | GGCACCCTCAGGTCATTCAT |
| *FOSB* | TCTGTCTTCGGTGGACTCCTTC | GTTGCACAAGCCACTGGAGGTC |
| *FOXL2* | GAGAAGAGGCTCACGCTGTC | GGTCCAGCGTCCAGTAGTTG |
| *HPRT* | TGACCTTGATTTATTTTGCATACC | CGAGCAAGACGTTCAGTCCT |
| *IL6* | AAAGAGGCACTGGCAGAAAA | TTTCACCAGGCAAGTCTCCT |
| **Mouse primers** | | |
| *Ak4* | TGGATTCACCCTTCTAGCGG | GTCTTAGCCTGGCAGCAACT |
| *Cox6a2* | GCCCTCTGCTCCCTTAACTG | GTGGGCAAAGGATTGACGTG |
| *Cyp19a1* | TACTTCATGTTACTTCTCGTCGC | TATCCTCGATCTTTATGTCTCTGTCAC |
| *Foxl2* | ACAACACCGGAGAAACCAGAC | CGAAGAACGGGAACTTGGCTA |
| *Hprt* | AGGACCTCTCGAAGTGT | ATTCAAATCCCTGAAGTA CTCAT |
| *Parp14* | GGTCTCTGTTGATGTGGGCACA | TTCTCGGTGGTCTTCAGGCAGT |
| *Thbs1* | CTCCAGCTCAGCTACCAACG | GGCCACAGATAGCTTGGAGG |
| *Xaf1* | CTGCGCTTCATAGTCCTTTGCC | AGGGTGCTGTTGGCTTTCCTTG |

**Supplementary Table 3 (large Excel file):** RNA-seq data showing differentially expressed genes in KGN, COV434 and AT29 cells

**Supplementary Table 4:** Genes of the MitoCarta3.0 database altered by EX-527 treatment in the three cell lines. Shown is the fold change (FC) induced by EX-527 in mouse (AT29) and human (KGN, COV434) GCT cells

| Gene name | Description | FC |
|---|---|---|
| **AT29 cells** | | |
| *Ak4* | adenylate kinase 4 | -7.71 |
| *Aldh1l2* | aldehyde dehydrogenase 1 family member L2 | 1.41 |
| *Bnip3* | BCL2 interacting protein 3 | -1.67 |
| *Bola1* | bolA family member 1 | -1.28 |
| *Chchd1* | coiled-coil-helix-coiled-coil-helix domain containing 1 | -1.34 |
| *Cmpk2* | cytidine/uridine monophosphate kinase 2 | 1.66 |
| *Cox6a2* | cytochrome c oxidase subunit 6A2 | 2.11 |
| *Dhcr24* | 24-dehydrocholesterol reductase | -1.30 |
| *Fam162a* | family with sequence similarity 162 member A | -1.66 |
| *Fam213a* | peroxiredoxin like 2A-PRXL2A | 1.27 |
| *Fdps* | farnesyl diphosphate synthase | -1.28 |
| *Gatm* | glycine amidinotransferase | -1.28 |
| *Gls2* | glutaminase 2 | 1.48 |
| *Idi1* | isopentenyl-diphosphate delta isomerase 1 | -1.34 |
| *Ldhb* | lactate dehydrogenase B | -1.29 |
| *Me3* | malic enzyme 3 | 1.34 |
| *Mgarp* | mitochondria localized glutamic acid rich protein | -1.59 |
| *Pdk1* | pyruvate dehydrogenase kinase 1 | -1.53 |
| *Rab32* | RAB32, member RAS oncogene family | -1.29 |
| *Tubb3* | tubulin beta 3 class III | -1.94 |
| **COV434 cells** | | |
| *ABAT* | 4-aminobutyrate aminotransferase | 1.35 |
| *ADCK1* | aarF domain containing kinase 1 | 1.32 |
| *ALDH6A1* | aldehyde dehydrogenase 6 family member A1 | 1.41 |
| *C15orf48* | chromosome 15 open reading frame 48-Cytochrome C Oxidase Subunit FA4 Like 3 | 1.31 |
| *CDC25C* | Cell Division Cycle 25C | -1.39 |
| *COQ3* | coenzyme Q3, methyltransferase | -1.36 |
| *DHODH* | dihydroorotate dehydrogenase (quinone) | -1.29 |
| *DTYMK* | deoxythymidylate kinase | -1.26 |
| *PIF1* | C15orf20\|PIF | -1.27 |
| *PRELID2* | PRELI domain containing 2 | -1.31 |
| *TIMM21* | translocase of inner mitochondrial membrane 21 | -1.28 |
| *TUBB3* | tubulin beta 3 class III | 1.34 |
| **KGN cells** | | |

#### Supplementary Table 4: continued

| | | |
|---|---|---|
| *AK4* | adenylate kinase 4 | 1.51 |
| *ALDH1B1* | aldehyde dehydrogenase 1 family member B1 | -1.28 |
| *BNIP3* | BCL2 interacting protein 3 | 1.49 |
| *COX18* | cytochrome c oxidase assembly factor COX18 | -1.26 |
| *FAM162A* | family with sequence similarity 162 member A | 1.34 |
| *FDPS* | farnesyl diphosphate synthase | -1.32 |
| *FTH1* | ferritin heavy chain 1 | 1.28 |
| *HK2* | sphingosine kinase 2 | 1.33 |
| *SLC25A36* | solute carrier family 25 member 36 | 1.48 |
| *THNSL1* | threonine synthase like 1 | 1.25 |
| *SDSL* | serine dehydratase like | 1.4 |
| *SUGCT* | succinyl-CoA:glutarate-CoA transferase | -1.29 |

**Supplementary Table 5:** Nrf2 target genes and ferroptosis-related genes in the three cell lines. Shown is the fold change (FC) induced by EX-527.

| **Gene name** | **Description** | **FC** |
|---|---|---|
| **Nrf2 targets** | | |
| **AT29 cells** | | |
| *Dusp5* | dual specificity phosphatase 5 | -1.28 |
| *Thbs1* | thrombospondin 1 | 1.30 |
| *Arrdc3* | Arrestin Domain Containing 3 | 1.80 |
| **KGN cells** | | |
| *ARRDC3* | arrestin domain containing 3 | 1.81 |
| *GDF15* | growth differentiation factor 15 | 1.35 |
| *FTH1* | ferritin heavy chain 1 | 1.28 |
| *HMOX1* | heme oxygenase 1 | 1.52 |
| **COV434 cells** | | |
| *FAM76B* | family with sequence similarity 76 member B | -1.28 |
| *FTL* | ferritin light chain | 1.46 |
| *ARRDC3* | arrestin domain containing 3 | 1.36 |
| **Ferroptosis (FerrDb)** | | |
| **AT29 cells** | | |
| *Mmp13* | Matrix Metallopeptidase 13 | 2.04 |
| *Arrdc3* | Arrestin Domain Containing 3 | 1.80 |
| *Il33* | Interleukin-33 | -1.52 |
| *Ddit3* | DNA Damage Inducible Transcript 3 | 1.53 |
| *Slc2a1* | Solute Carrier Family 2 Member 1 | -1.61 |
| *Bnip3* | BCL2 Interacting Protein 3 | -1.67 |
| **KGN cells** | | |
| *HIC1* | Hypermethylated In Cancer 1 | -1.60 |
| *ARRDC3* | Arrestin Domain Containing 3 | 1.81 |
| *IL6* | Interleukin-*6* | 1.78 |
| *TFRC* | Transferrin Receptor | 1.54 |
| *HMOX1* | Heme Oxygenase-1 | 1.52 |
| *BNIP3* | BCL2 Interacting Protein 3 | 1.5 |

#### Supplementary Table 6: Individual AT83 mouse tumor volumes during EX-527 treatment.

AT83 mouse tumor volumes (in $mm^3$) were measured using Vevo LAZR high-frequency ultrasonic-photoacoustic imager each week, in vehicle-treated (control, CTR) and EX-527-treated groups. R: right ovary, L: left ovary. Tumor volumes of the #507 mouse were not assessed during the last week due to death before the end of treatment.

| | | Treatment time ( weeks) | | | | | |
|---|---|---|---|---|---|---|---|
| | **Tumors #** | **0** | **1** | **2** | **3** | **4** | **Gain of volume** |
| **CTR** | 469 R | 320.0 | 402.6 | 482.0 | 469.0 | 483.0 | + |
| | 469 L | 583.0 | 589.9 | 656.8 | 768.4 | 772.0 | + |
| | 474 R | 375.5 | 336.7 | 258.3 | 262.8 | 487.1 | + |
| | 474 L | 701.4 | 963.3 | 1071.0 | 1565.0 | 1620.0 | + |
| | 500 R | 1960.5 | 1837.3 | 2399.0 | 2874.0 | 2843.3 | + |
| | 516 R | 59.0 | 45.6 | 59.0 | 70.0 | 145.5 | + |
| | 516 L | 113.5 | 115.9 | 175.0 | 247.9 | 490.5 | + |
| **EX-527** | 517 R | 70.9 | 157.1 | 247.0 | 265.0 | 202.0 | + |
| | 517 L | 205.9 | 280.7 | 293.0 | 297.3 | 447.2 | + |
| | 518 R | 27.0 | 18.0 | 24.5 | 31.8 | 54.3 | + |
| | 518 L | 16.6 | 17.0 | 33.0 | 21.6 | 21.8 | = |
| | 462 R | 2039.0 | 1645.2 | 1286.0 | 1323.0 | 1766.0 | - |
| | 462 L | 2973.6 | 2268.3 | 2051.0 | 2470.4 | 2919.0 | = |
| | 487 R | 2216.9 | 2846.8 | 3448.0 | 2505.0 | 2123.0 | - |
| | 487 L | 1209.5 | 956.6 | 1255.0 | 830.0 | 1253.0 | = |
| | 507 R | 227.3 | 165.2 | 116.2 | 118.6 | / | - |
| | 507 L | 1979.8 | 1453.4 | 1405.1 | 1644.0 | / | - |
